# Effects of D3-preferring agonist PD 128907 on compulsive behaviour and decision making as assessed by the 5C-CPT

**DOI:** 10.1101/2024.01.22.576339

**Authors:** Sara Abdulkader, John Gigg

## Abstract

**Background:** Repetitive rituals in OCD patients result from pathological doubt, which has been linked to dysfunction in decision-making. First-line drug treatments for OCD are selective serotonin reuptake inhibitors; however, 40% of OCD patients do not respond to these. As brain activity patterns in OCD resemble those in schizophrenic patients, this suggests a dopaminergic component, supported by data showing that the D2/D3 receptor agonist Quinpirole induces compulsive checking in male rats. OCD has proved difficult to model in rodents and the contribution of decision making to compulsive behaviour in such models has not been studied. The five-choice continuous performance task (5C-CPT) measures both decision making and compulsive behaviour, making it possible to evaluate whether OCD models show correlated changes in these two behaviours. Establishing this would provide a new model approach to help develop therapeutic agents for OCD.

**Aims:** The role of dopaminergic D3 receptors in decision making and compulsive behaviour was determined by testing the effect of the preferential D3 agonist PD 128907 on 5C-CPT performance measures. Oldham’s method was used to determine the presence of any rate-dependent effect.

**Methods:** Female Lister hooded rats were trained to criterion in the 5C-CPT (>70% accuracy, < 30% omission and < 40% false alarms). The effects of PD128907 (0.25-1 mg/kg) were then investigated under challenging task conditions.

**Results:** Oldham method revealed: a moderate positive association between baseline perseverative correct responses and the change at 0.05 mg/kg PD 128907; a strong positive association between baseline perseverative false alarm and the change at 0.2 mg/kg; a positive association between baseline total number of perseverative responses and the change at 0.05 mg/kg or 0.2 mg/kg PD 128907; a positive relationship between baseline accuracy and the change at 0.05 mg/kg PD 128907; and a moderate positive association between baseline correct response latency and the change at 0.05 mg/kg or 0.2 mg/kg PD 128907. The angle measurements and the direction of movement (clockwise or counterclockwise) showed how effective is one dose at increasing compulsive behaviour compared with other doses.

**Conclusions:** PD 128907 effects on compulsive like behaviour and decision making in poor performing female lister hooded rats with long correct response latency and high perseveration at baseline (vulnerable population) in the 5C-CPT task matches two key features of OCD compulsions in humans (perseveration and indecision), this suggests that PD 128907 is more effective than Quinpirole in simulating the brain network conditions that underpin OCD. This model could help to develop more successful pharmacological interventions and to generate data translatable to clinical studies.

## Introduction

Compulsive behaviour is present in different neuropsychiatric disorder such as obsessive-compulsive disorder (OCD), compulsive drug seeking behaviour, compulsive gambling and schizophrenia (Nestadt et al., 2016). Obsessive-compulsive disorder is a neuropsychiatric condition characterized by obsessions (recurrent thoughts) that are neutralised by compulsions (repetitive motor actions) (Miodownik et al., 2015). These symptoms greatly affect social and emotional functioning (Asnaani et al., 2017). Pathological doubt is recognized as a core problem in OCD and it has been linked to impairment in decision making process (Nestadt et al., 2016). The prevalence of OCD is between 1-3%, however, OCD is often unrecognised and undertreated (Mathes et al., 2019). OCD is more prevalent among females in adolescence and adulthood compared to males (Mathes et al., 2019). Depending on the main symptoms, OCD can be divided into three categories: predominantly obsessional thoughts, predominantly compulsive acts and a third subtype that is a combination of the first two (van Oudheusden et al., 2020). OCD symptoms are linked with anxiety and may lead to social isolation (Hillman et al., 2022). Selective serotonin reuptake inhibitors (SSRI) are the first-line pharmacological treatment for OCD (Goodman et al., 2021). However, 40% of patients with OCD do not respond to current medications and individuals with OCD have lower quality of life, even after treatment, compared with healthy individuals (Miodownik et al., 2015; Jahangard et al., 2018). Therefore, there is need for new therapeutic agents.

Dopamine receptors are divided into two families: D1-like and D2-like receptors (including D2, D3 and D4 receptors) (Nakajima et al., 2013). D3 receptors are involved in social behaviour, motor and cognitive functions (Nakajima et al., 2013). They serve as both post-synaptic receptors and autoreceptors. They are mainly located in regions of the limbic system, including the nucleus accumbens, but are also found in the frontal and other cortical regions (Herroelen., et al, 1994; Bouthenet et al. 1991). In addition, D3 receptors have the highest affinity for dopamine (Kehr et al., 2022). Despite its unique properties there is no selective D3 receptor agonist (Watson et al., 2012). D3 *preferring* agonists have been used to explore the role of D3 in cognitive function (Millan et al., 2010; Watson et al., 2012). For example, PD128907 is a D3 receptor preferring agonist that has a 54-fold preference for D3 over D2 receptors (Pugsley et al., 1995). PD128907 impaired performance in rats for novel object recognition and social novelty discrimination (Watson et al., 2012). Conversely, the D3 receptor preferring antagonist S33138, which has a 25-fold preference for D3 over D2 receptors, improved performance in the same tasks (Millan et al., 2010). According to these studies, D3 preferring antagonists can be used for treatment of schizophrenia and are not expected to elicit motor side effects (extrapyramidal side effects) classically elicited by D2 receptor antagonists (Watson et al., 2012). Two D3 receptor preferring D3R/D2R antagonists reached phase II clinical trials for treatment of schizophrenia (Kiss et al., 2021). F 17464 which has 38-71-fold preference for D3 over D2 receptor caused a significant improvement in patient with acute exacerbation of schizophrenia (Bitter et al., 2019).

There is an elevated risk of developing substance abuse in subjects with existing OCD (Rowe et al., 2022). D3 receptors are less abundant than D2 receptors. In addition, the nucleus accumbens (associated with compulsive behaviour) has a very high D3 receptor density, while D3 receptor density in dorsal striatum (associated with motor function) is very low (Sokoloff and Le Foll, 2017). This receptor distribution in the brain links it with compulsive drug-seeking behaviour, for example, studies suggest that D3 receptors are involved in cocaine’s action and D3 receptor agonism enhances cocaine seeking behaviour (Xi et al., 2004). Conversely, D3 receptor antagonism suppresses drug seeking behaviour, suggesting that D3 receptors can be used as a target for developing anti-addiction medications (Vorel et al., 2002). Furthermore, the D3 receptor has also been associated with gambling (Zeeb et al., 2009). OCD demonstrate high comorbidity with compulsive gambling (Medeiros et al, 2018). OCD and pathological gambling are part of behavioural addiction. In the rodent gambling task (based in part on the Iowa gambling task for human subjects) dopaminergic agents appear to modulate gambling behaviour; for example, the non-selective D2/3/4 antagonist Eticlopride improved gambling-related decision making without affecting waiting impulsivity in this task (Zeeb et al., 2009). However, the more selective D3 receptor agents such as PD128907 did not affect decision making and motor impulsivity in the same task (Di Ciano et al., 2015).

Studies employing 5-CSRTT to determine the effect of the D3 receptor in modulating cognitive function and inhibitory response control have produced conflicting results. For example, some studies report that D3 receptor agonism impairs accuracy (Zhu et al., 2017), while others report that D3 receptor blockade has no effect on performance (% correct response and omission) (Millan et al., 2008). On the other hand, when it comes to inhibitory response control, D3 receptor agonism appears to improve waiting impulsivity; however, this effect could have been due to a decrease in motivational performance (Zhu et al., 2017). Other studies report that D3 receptor blockade by S33138 does not affect waiting impulsivity in the 5-CSRTT (Millan et al., 2008). It would be insightful to investigate effect of PD 128907 on a task with higher translational validity to human such as 5C-CPT. 5C-CPT is a valuable tool for assessment of several cognitive domains such as attention, decision making and compulsive behaviour (Barnes et al., 2012; Tomlinson et al., 2014). Dysfunction in all these cognitive domains has been reported in OCD patients (Benzina et al., 2016). D3 receptors are expressed predominantly in the mesolimbic pathway, which has been associated with impulsive behaviour, compulsive behaviour, and addiction (Herroelen., et al, 1994; Bouthenet et al. 1991). The role of PD128907 in modulating cognitive function and inhibitory response control in the 5C-CPT remains unknown. Understanding the contribution of D3 receptors to attention, impulsive and compulsive behaviour as measured by 5C-CPT will help the development of selective therapeutic agents for OCD, compulsive drug seeking behaviour, ADHD, compulsive gambling and cognitive deficit in schizophrenia. In addition, Alteration of 5C-CPT performance measures by systemic administration of D3 preferring agonists may present a valuable tool for understanding the neurobiological basis of OCD and other compulsive behaviour related disorders. The purpose of the present study is to investigate the contribution of D3 receptor on the performance of female Lister hooded rats in the 5C-CPT task via systemic administration of D3 preferring agonist PD 128907. Correlational analysis may present an opportunity to enhance the translational value of research in the preclinical laboratory to the clinic (Abdulkader et al., 2023a; 2023b). Most studies examining effects of D2/D3 receptor agonist (such as Quinpirole) on compulsive behaviour has ignored individual variability in baseline performance by using average performance of all rats. Previous studies report that compulsive behaviour and waiting impulsivity change rate dependently in the 5C-CPT following administration of dopaminergic agents such as GBR 12909, SKF 38393 or amphetamine (Abdulkader et al., 2023a; 2023b). Therefore, Oldham’s correlative approach was used to determine if PD 128907 can modulate 5C-CPT performance measures in a rate-dependent manner. We hypothesised that PD 128907 would change inhibitory response control and cognitive function in a manner consistent with baseline performance.

## Materials and Methods

### Animals

Female Lister hooded rats (n=15; Charles River, UK) were housed in groups of 5 in a temperature (21 ± 2 °C) and humidity (55±5%) controlled environment (University of Manchester BSF facility). Rats weighed 230±10g upon arrival. They were allowed free access to food (Special Diet Services, UK) and water for one week prior to the beginning of training. Home cages were individually ventilated cages with two levels (GR1800 Double-Decker Cage, Techniplast, UK) and testing was completed under a standard 12-hour light: dark cycle (lights on at 7:00 am). Two days before training commenced food restriction was initiated to encourage their task engagement and this continued throughout training. Rats were maintained at approximately 90% of their free-feeding body weight (fed 10 g rat chow/rat/day) and were provided with free access to water. All procedures were conducted in accordance with the UK animals (Scientific Procedures) 1986 Act and local University ethical guidelines.

### Rat 5C-CPT Apparatus

Training took place in 8 operant chambers (Campden instruments Ltd, UK), each placed inside a ventilated and sound-attenuating box. The chambers were of aluminium construction 25cm × 25 cm. Each chamber had a curved wall containing nine apertures. For 5C-CPT training, four of the apertures were blocked by metal caps while the other five were left open (numbers 1, 3, 5, 7 and 9 were open). Each aperture had a light and an infrared beam crossing its entrance to record beam breaks following nose poke. A reward dispenser was located outside each chamber that automatically dispensed reinforcers (45 mg sucrose food pellets; Rodent Pellet, Sandown Scientific, UK) into a food tray at the front of the chamber. Entrance to each food tray was covered by a hinged panel so that tray entries were recorded when the rat’s snout opened the panel. House lights were placed in the side wall of the chambers and performance was monitored by a camera attached to the roof.

### 5C-CPT behavioural training

The 5C-CPT is a modified version of the 5-CSRTT and incorporates non-target trials in which all five holes are lit and the rat must not respond at any of the five positions to be rewarded (Young et al., 2009). The incorporation of non-target trials helps to dissociate response inhibition from waiting impulsivity; premature responses during target trials reflect a deficit in waiting impulsivity, while the failure to withhold responding (false alarms) during non-target trials represents response disinhibition (Young et al., 2011). This task also allows the assessment of vigilance in rodents in a manner consistent with human CPT using signal detection theory (Barnes et al., 2012; Tomlinson et al., 2014). In addition, a variable ITI is incorporated into the 5C-CPT to prevent rats from simply timing their response (Van Enkhuizen et al., 2014). Training was conducted in a similar manner to previous studies using 5C-CPT (Barnes et al., 2012; Tomlinson et al., 2014) with training carried out 5 days per week. A session consisted of 120 trials or was 30 min long, whichever came first. The training consisted of 3 phases: (i) 5-CSRTT training, (ii) 5C-CPT training with fixed ITT and (iii) 5C-CPT training with variable ITI (Barnes et al., 2012; Tomlinson et al., 2014). Task parameters such as stimulus duration and limited hold were gradually reduced for rats based on their individual performance. It took 4 months for the animals to discriminate between target and non-target trials and one month for animals to achieve a stable baseline performance. The main variables of interest were number of perseverative responses, response latency and accuracy (Table 1)

**Table 1:** Description of 5-CPT outcome measures.

| **Outcome measures** | **Descriptions** | **Dimension** |
| --- | --- | --- |
| **Perseverative correct response** | Repeated nose pokes after making a correct response | Compulsive behaviour |
| **Perseverative false alarm** | Repeated nose poked after making a false alarm | Compulsive behaviour |
| **Total number of Perseverative responses** | Repetitive responses (repeated nose pokes) even after making a response | Compulsive behaviour |
| **Correct latency** | Time taken to make a correct response | Decision making |
| **Accuracy** | Proportion of correct responses to incorrect response | Selective attention |

### Signal detection theory (SDT)

Two primary parameters are calculated from 5C-CPT trial-outcome measures. The first is hit rate, which is the proportion of ‘hits’ to ‘misses’ and the second is false alarm rate, which is the proportion of false alarms to correct rejections (Young et al., 2011). From these two parameters, sensitivity (vigilance) and bias can be calculated using SDT (see supplement). Sensitivity reflects the ability to respond to the illuminated hole and inhibit that prepotent response when all five holes are illuminated (Barnes et al., 2012; Tomlinson et al., 2014; Hayward et al., 2016) Sensitivity can be estimated by calculating the discriminability index (*d*’), which is the difference between z score of hit rate and false alarm rate (Young et al., 2009; Anderson, 2015). 5C-CPT performance can also be affected by bias (tendency to respond to stimuli), which can be measured by calculating the responsivity index (Barnes et al., 2012; Tomlinson et al., 2014; Hayward et al., 2016). A positive value for RI reflects bias towards FA (liberal response strategy), while a negative RI reflects bias towards omission (conservative response strategy). When RI is zero there is no bias (Anderson, 2015).

### Experimental Design

#### Drug preparation

D3 receptor agonist PD128907 (Sigma-Aldrich) were dissolved in 0.9% saline to obtain the required final concentration. The doses were based on previous reports (Di Ciano et al., 2015; Zhu et al., 2017). The drug was injected via the intraperitoneal (i.p.) route 10 min before behavioural testing at a volume of 1 ml/kg. All drug doses were calculated as base equivalent weight.

### Behavioural testing

Fifteen rats were used to investigate the effect of PD128907 on 5C-CPT performance using challenging task conditions that consisted of a variable ITI (8, 10 and 12 s). The session lasted no longer than 60 min. Using a within-subject design rats received 0.05, 0.1, 0.2 mg/kg PD128907 or vehicle. All animals received a normal training session between the testing sessions to ensure performance was maintained at a stable baseline. Each test day was followed by at least a 7-day washout period, after which behavioural testing commenced. Prior to the first test day, all animals had been habituated twice to i.p. saline injections.

### Statistical analysis

The main variables of interest were number of perseverative responses, response latency and accuracy. All data were displayed as observed mean ± SEM. Overall performance was analysed by one-way repeated measures ANOVA followed by Planned Comparisons. Analysis of the performance across the prolonged ITI was carried out by two-way repeated measures ANOVA (ITI and Treatment as within-subject factors), followed by Planned Comparisons. Alpha level was set to 0.05 and all analysis was carried out using Prism (v10.0).

### Correlational analysis

This study investigated if baseline performance predicted the degree of change in performance following administration of PD 128907 using Oldham method. Oldham’s method was used to determine the presence of a rate dependent relationship. Pearson’s correlation was used when individuals were normally distributed. Nonparametric Spearman’s correlation was used when individuals were not normally distributed. When interpreting the output from Oldham’s technique the following conclusions can be reached: *r*oldham > 0.3 demonstrates the presence of a rate-dependent relationship; *r*oldham = 0.3-0.6 demonstrates the presence of a moderate relationship between baseline performance and the change; and *r*oldham > 0.7 demonstrates the presence of a strong relationship between baseline performance and the change (Oldham, 1962; Snider et al., 2016; Bickel et al., 2016; Abdulkader et al., 2023a). A linear regression was fitted to enhance visual interpretation of the data.

## Results

### Compulsive behaviour

The main effect of PD128907 on compulsive behaviour is shown in Fig. 1. PD128907 showed a tendency toward increasing perseverative responding during target trials (perseverative correct response) at 0.05 mg/kg, although this did not reach significance (F (1.964, 25.54) = 1.283, P=0.29; Fig 1A). Oldham method revealed a moderate positive association between baseline perseverative correct responses and the change following administration of the lowest dose of PD 128907, indicative of a rate dependent effect (Fig. 1B). The moderate positive correlation changed to zero following administration of the moderate dose of PD128907 (0.58 vs 0.00) and it switched to an insignificant negative relationship following administration of the highest dose of PD 128907 (0.58 vs -0.36). In other words, the strength and the type of the relationship gradually changed as the dose increased. PD 128907 changed perseverative correct responses in a dose dependent. Oldham’s angle measurements for the relationship between baseline perseverative correct responses and PD 128907 induced change are shown in Fig 1B. Amount of space between the regression lines were measured using a protractor. Three angles were measured: ∠1, ∠2 and ∠3 as shown in Fig 1B. ∠1 is the space between 0.05 mg/kg and 0.2 mg/kg regression lines. ∠2 is the space between 0.1 mg/kg and 0.2 mg/kg regression lines. ∠3 is the space between 0.05 mg/kg and 0.1 mg/kg lines The measurement of the angle and the direction of movement (clockwise or counterclockwise) shows how effective is one dose at increasing compulsive behaviour compared with other doses.

**Figure 1:**
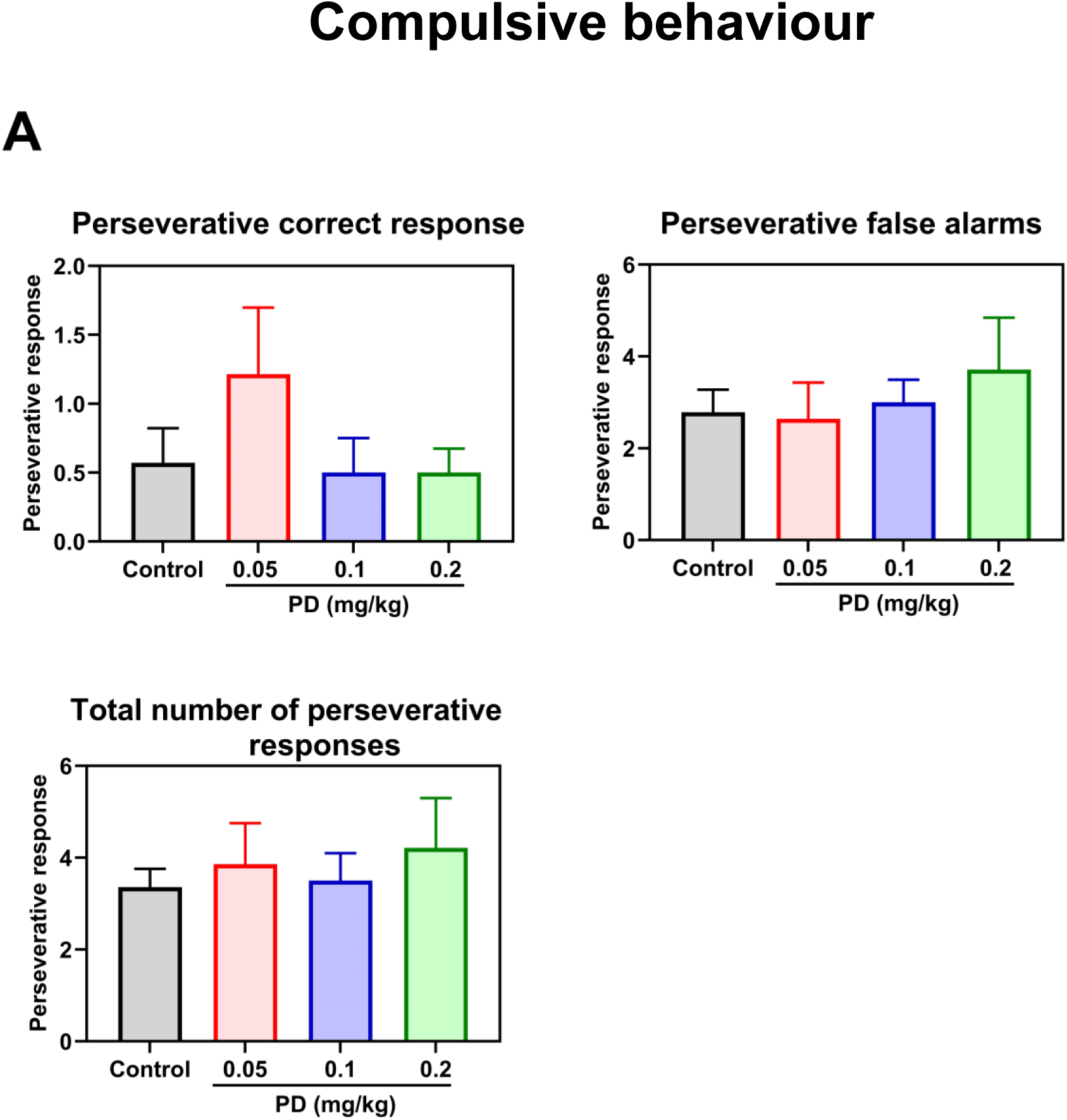
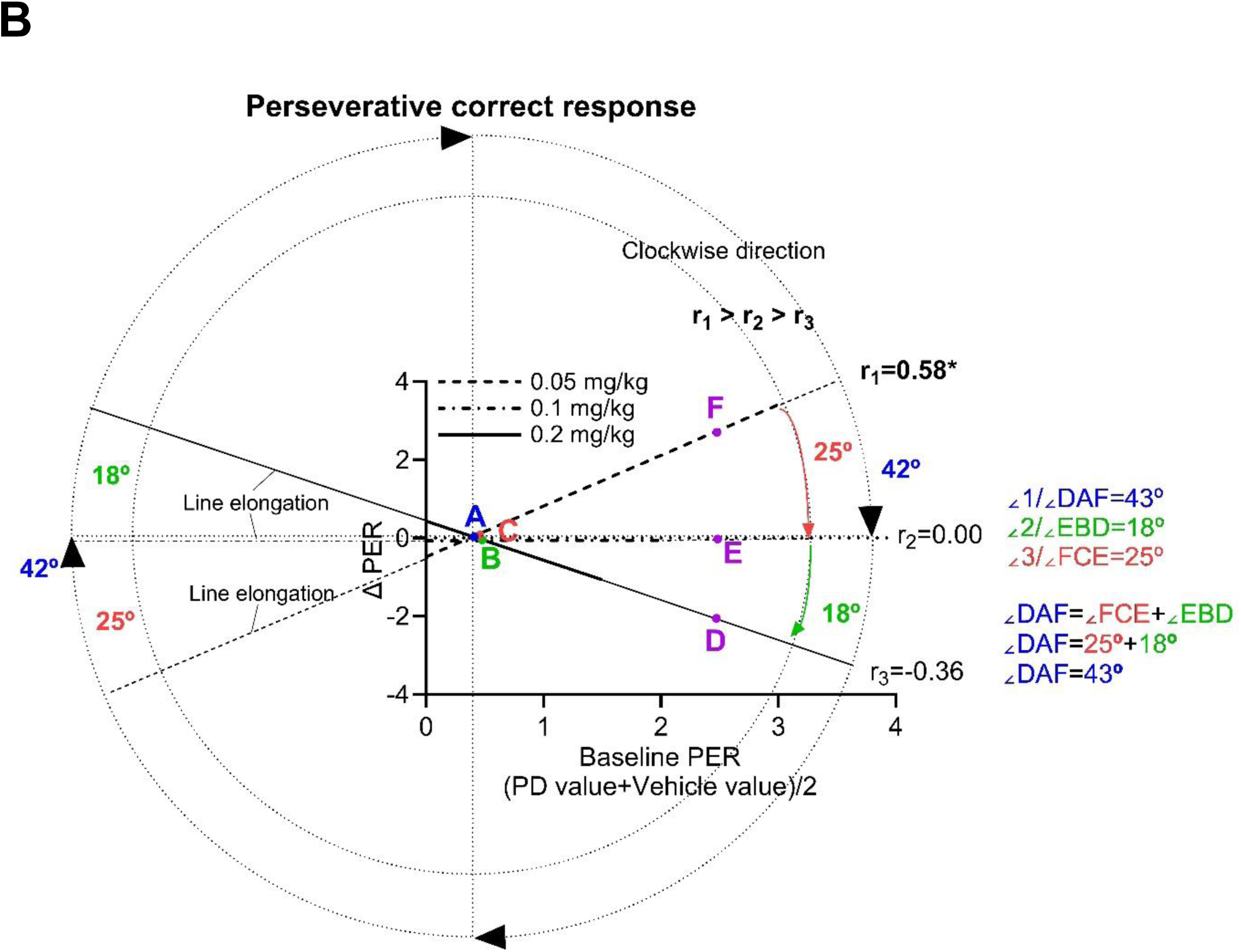
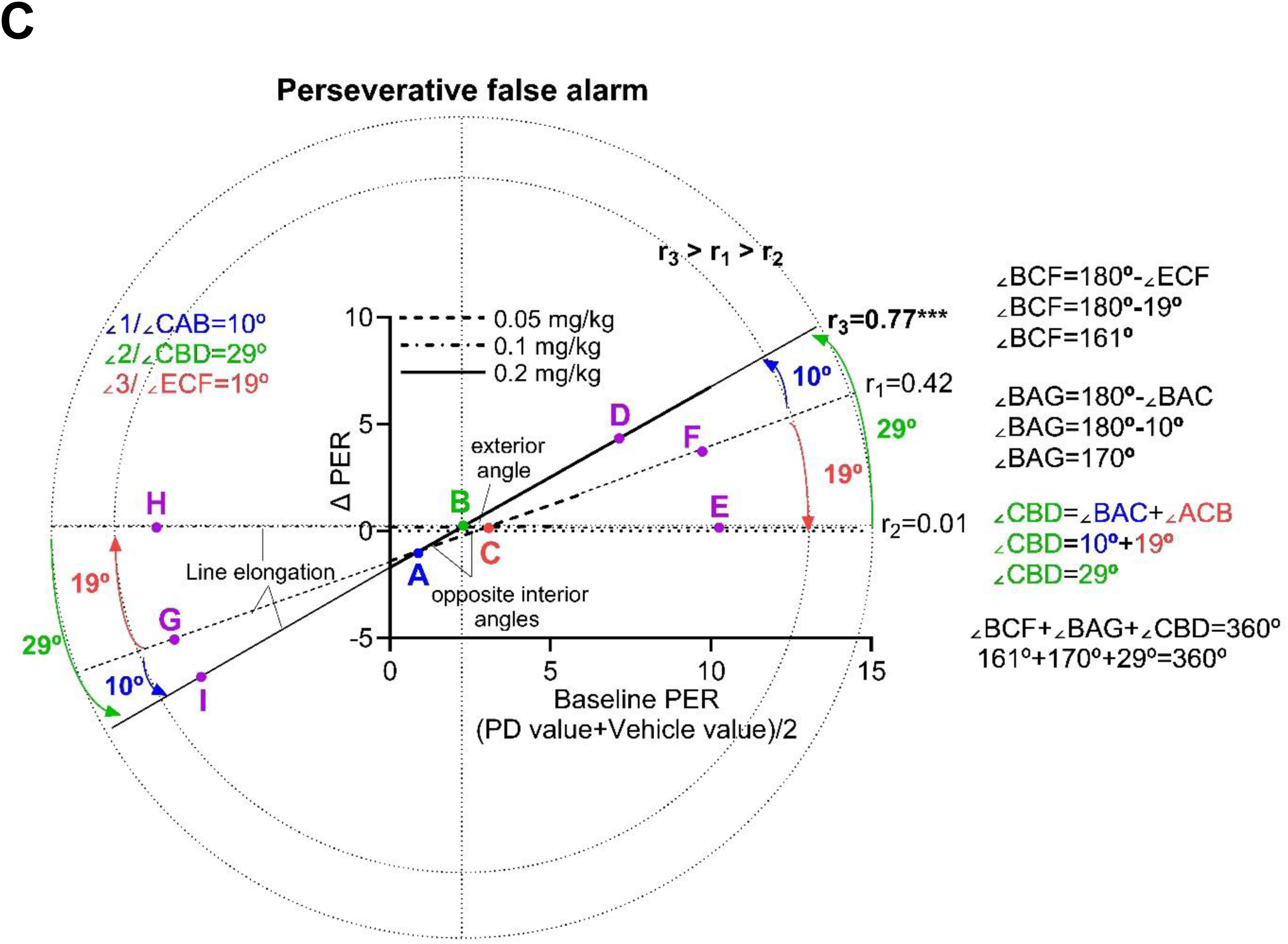
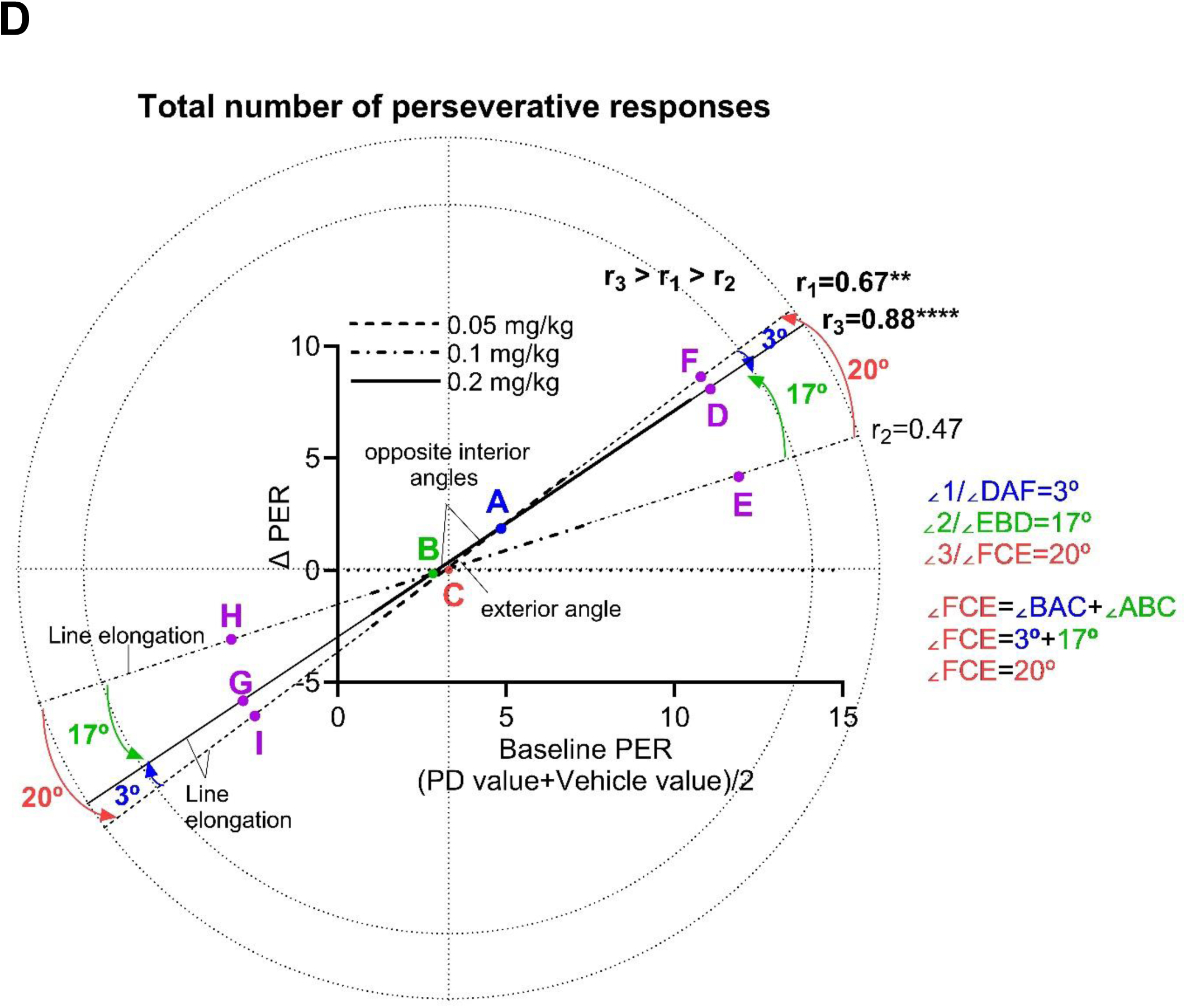
Effects of PD128907 (0.05, 0.1 or 0.2 mg/kg) on perseverative responding in the 5C-CPT. (A) Effects of PD128907 on perseverative correct response, perseverative false alarms, and total number of perseverative responses. (B) Relationship between baseline perseverative correct responses and PD 128907 induced change. (C) Relationship between baseline perseverative false alarms and PD 128907 induced change. (D) Relationship between baseline total number of perseverative responses and PD 128907 induced change. Arrows show the clockwise or counterclockwise movement of the regression line as the dose increases. PER= Perseverative response.

PD128907 showed a tendency toward increasing perseverative responding during non-target trials (perseverative false alarm) at 0.2 mg/kg (F (F (2.070, 26.90) = 0.7417, P=0.49; Fig. 1A). Oldham method revealed a strong positive association between baseline perseverative false alarm and the change at 0.2 mg/kg, indicative of a rate-dependent effect (Fig 1C). The counterclockwise movement of the regression line shows that, when the dose increases, the effectiveness of the drug increases. Oldham method revealed an insignificant moderate positive relationship between baseline perseverative false alarm and the change following administration of the lowest dose of PD 128907. This relationship changed to a significant strong positive correlation following administration of the highest dose of PD 128907 (0.42 vs 0.77). The strength of the relationship increased as the dose increased. However, the intermediate dose was less effective than the lowest dose at increasing the number of perseverative false alarms. This could be due to change in bias (tendency to respond to stimuli). Oldham’s angle measurements for the relationship between baseline perseverative false alarms and PD 128907 induced change are shown in Fig 1C. ∠1 and ∠3 are two opposite interior angles. ∠2 is the exterior angle of the triangle. The Exterior Angle theorem states that the exterior angle of a triangle is equal to the sum of the two opposite interior angles. If two angles are known, then the exterior angle theorem can be used to calculate the third angle. For example, ∠2=∠1+∠3.

PD128907 produced no significant change in perseverative responding when the number of perseverative responses during target and non-target trials were combined (F (2.322, 30.18) = 0.4425, P = 0.67; Fig. 1A). PD128907 showed a tendency toward increasing total number of perseverative responses at 0.05 mg/kg or 0.2 mg/kg (Fig 1A). Oldham method revealed a strong positive association between baseline compulsivity (perseverative correct responses + perseverative false alarms) and the change at 0.05 and 0.2 mg/kg (Fig 1D). Rats with baseline compulsive behaviour > 5 impaired the most following the treatment. The strength of the relationship between baseline compulsive behaviour and PD 128907 induced change diminished at 0.1 mg/kg compared with 0.05 mg/kg. The change in total number of perseverative responses following administration of the lowest dose of PD128907 is mainly due to change in perseverative correct response while the change in total number of perseverative responses following administration of the highest dose of PD128907 is mainly due to the change in perseverative false alarm, suggesting that these two types of behaviour should be considered separately. Oldham’s angle measurements for the relationship between baseline total number of perseverative responses and PD 128907 induced change are shown in Fig 1D. ∠1 and ∠2 are two opposite interior angles. ∠3 is the exterior angle of the triangle. If two angles are known, then the third angle can be calculated. ∠3=∠1+∠2 (Fig 1D). It is hard to understand the movement of the regression line as the dose increases, suggesting that perseverative correct responses and perseverative false alarms should be considered separately.

### Selective attention and response latency

The main effect of PD128907 on accuracy and response latency is shown in Fig. 2. One-way repeated measures ANOVA revealed no significant treatment effect [F (2.11, 27.43) = 1.7, P = 0.198; Fig 2A] on accuracy. Oldham method revealed a significant positive relationship between baseline accuracy and the change following administration of 0.05 mg/kg PD 128907. Accuracy of rats with baseline accuracy < 95% impaired the most after the treatment with 0.05 mg/kg PD128907. The movement of the regression line in the clockwise direction shows that, when the dose increases, the positive relationship between baseline accuracy and PD 128907 induced change gradually disappears (decreased effectiveness). The significant positive relationship at the lowest dose of PD 128907 changed to an insignificant positive relationship following administration of the moderate dose of PD 128907 (0.62 vs 0.46) and the relationship disappeared entirely at the lowest dose of PD 128907 (0.62 vs 0.15). The strength of the relationship decreased gradually as the dose increased. PD 128907 changed accuracy in a dose dependent manner. Oldham’s angle measurements for the relationship between baseline accuracy and PD 128907 induced change are shown in Fig 2B. ∠2 and ∠3 are two opposite interior angles. ∠1 is the exterior angle of the triangle. The Exterior Angle theorem states that the exterior angle of a triangle is equal to the sum of the two opposite interior angles. If two angles are known, then the exterior angle theorem can be used to calculate the third angle. For example, ∠1=∠2+∠3.

**Figure 2:**
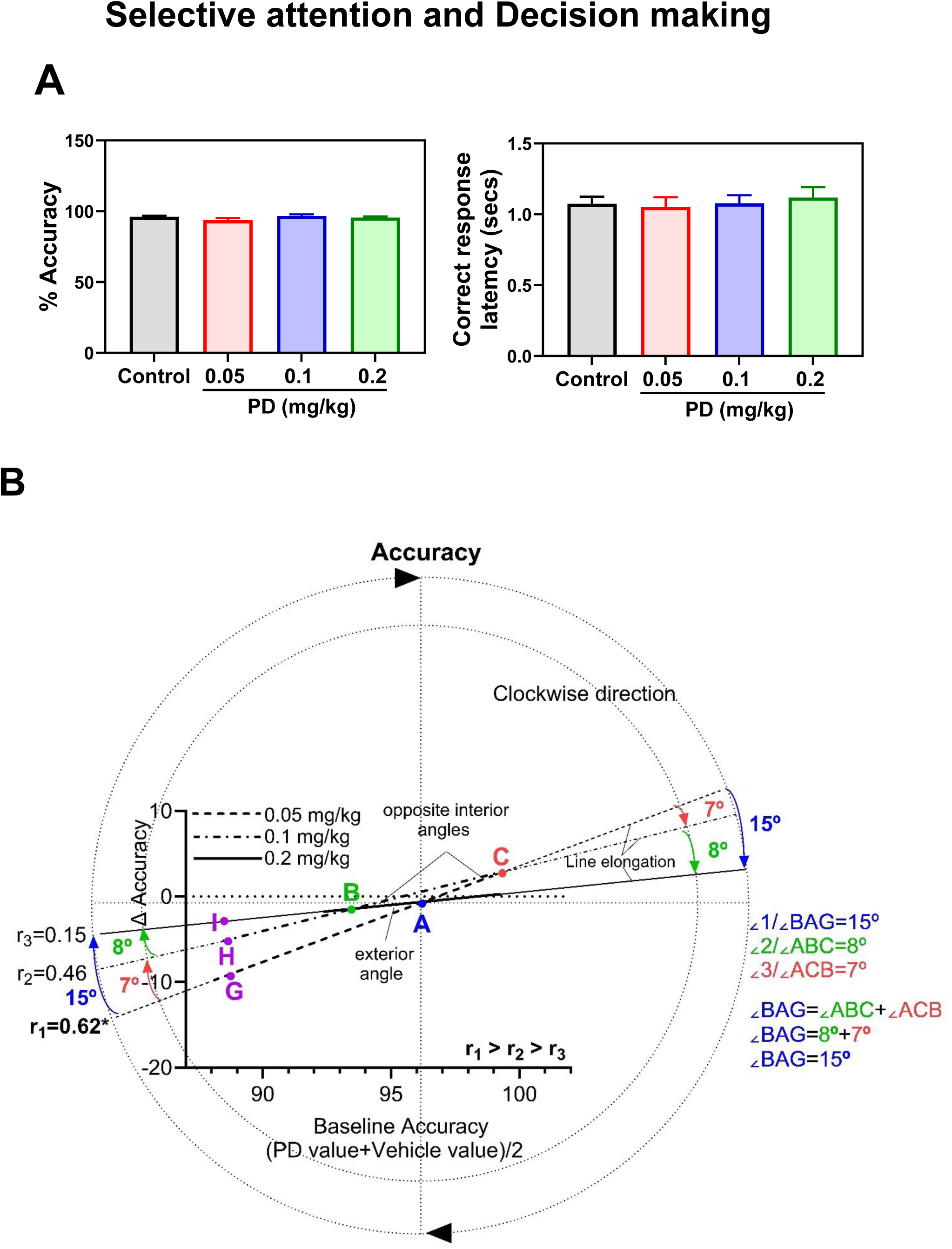
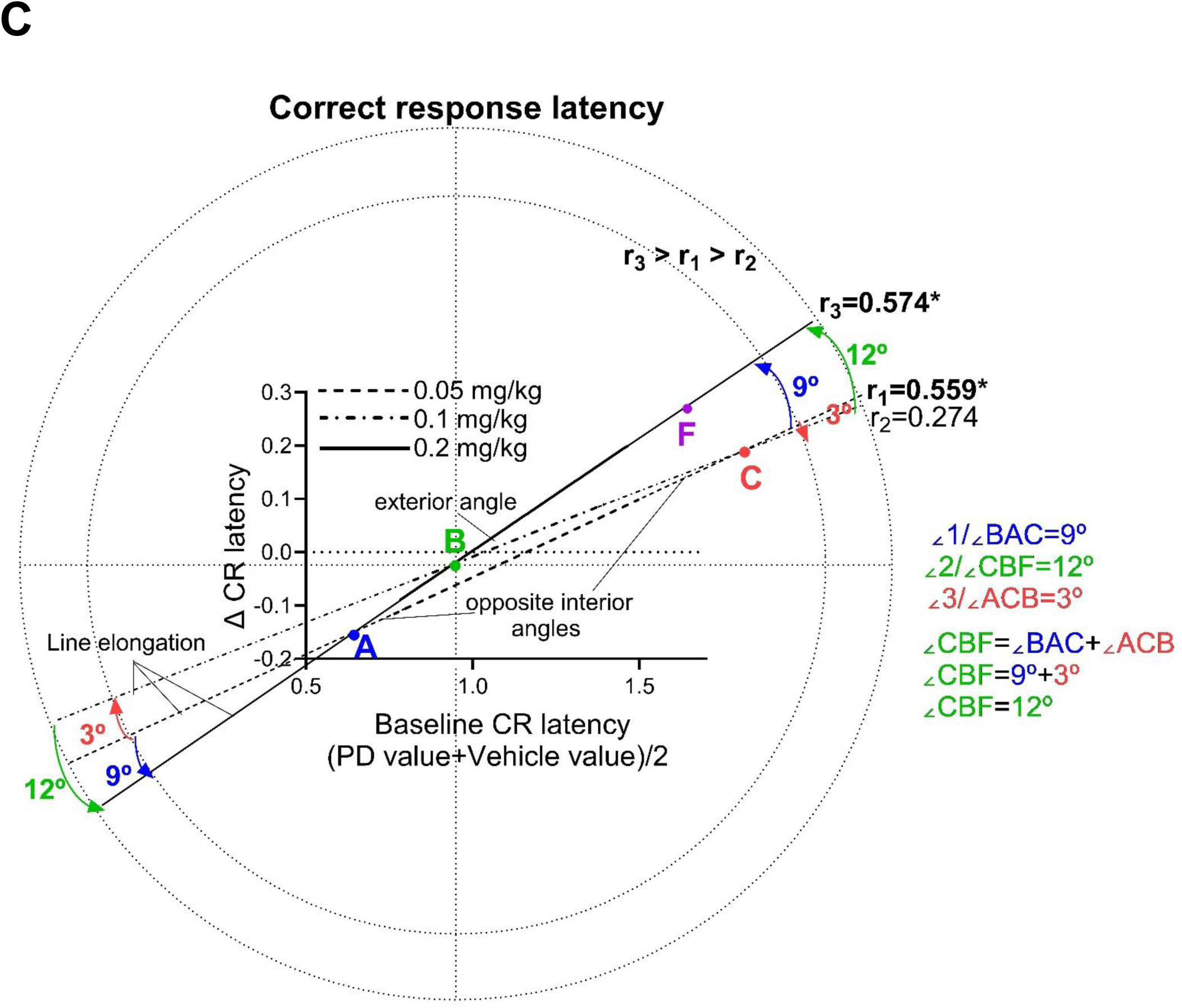
Effects of PD128907 (0.05, 0.1 or 0.2 mg/kg) on accuracy and correct response latency. (A) The traditional method of presenting the data. (B) Relationship between baseline accuracy and PD 128907 induced change. (C) Relationship between baseline correct response latency and PD 128907 induced change. The arrows show the clockwise or counterclockwise movement of the regression line as the dose increases. CR= correct response.

One-way repeated measures ANOVA revealed no significant treatment effect [F (2.408, 31.30) = 0.6989, P = 0.69; Fig 2A] on correct response latency. Consistent with effect of PD 128907 on compulsive behaviour, Oldham method revealed a significant positive association between baseline correct response latency and the change in correct response latency following administration of 0.05 mg/kg or 0.2 mg/kg PD 128907, indicative of a rate dependent effect (Fig 2C). The Oldham regression line moved in counterclockwise direction as the dose increased. The type of the relationship did not change as the dose increased. The significant positive relationship at the lowest dose of PD 128907 changed to an insignificant positive relationship following administration of the moderate dose of PD 128907 (0.559 vs 0.27). A stronger relationship was observed at the highest dose of PD 128907 compared with the lowest dose of PD 128907 (0.57 vs 0.559), indicative of increased effectiveness. Line elongation showed that ∠1 and ∠3 are two opposite interior angles. ∠2 is the exterior angle of the triangle (Fig 2C). If two angles are known, then the exterior angle theorem can be used to calculate the third angle. For example, ∠2=∠1+∠3.

## Discussion

The purpose of the present study was to determine the effects of the D3-preferring agonist PD128907 on 5C-CPT performance under challenging task conditions. The main finding of this experiment was that the lowest dose of PD128907 changed accuracy, correct response latency (decision making) and perseverative correct responses (compulsive behaviour) in a rate-dependent manner. On the other hand, the highest dose of PD128907 changed perseverative false alarms (compulsive behaviour) and correct response latency (decision making) in a rate-dependent manner. This study suggests that :1) D3 receptors make contribution to decision making and compulsive like behaviour (perseverative responses) in the 5C-CPT; 2) PD 128907 induced behavioural alterations (perseveration and indecision) in female lister hooded rats that have a high perseveration and a long correct response latency at baseline (vulnerable population) can be used as an animal model of OCD; 3) Change in decision making (increase in the time taken to make a response) and change in compulsivity (perseveration) co-occur; 4) the effect of PD 128907 on compulsive behaviour, decision making and attention is dependent on heterogeneity of the performance prior to manipulation.

Clinical studies suggest that poor decision-making contributes to OCD symptoms in patients (Nestadt et al., 2016). In the 5C-CPT, increase in the time taken to make a correct response represents diminished certainty in reaching a decision (Barnes et al., 2012; Tomlinson et al., 2014). Dysfunction in mesocorticolimbic dopaminergic system has been linked with deficit in decision making (St Onge et al., 2011). In the present study, the lowest dose of PD 128907, which affected perseverative correct response, changed correct response latency in a rate-dependent manner. On the other hand, the highest dose of PD 128907, which changed perseverative false alarm, changed correct response latency in a rate dependent manner. This finding suggests that poor decision making (impairment in correct response latency) may have contributed to compulsive like behaviour (increased perseveration) in poor performing rats. Previous studies report that at doses lower than 0.1 mg/kg PD 128907 is D3 receptor selective (Zapata et al., 2001). These findings suggest that effect of 0.05 mg/kg PD 128907 on perseverative correct response could be due to D3R agonism while effect of 0.2 mg/kg PD 128907 on perseverative false alarm could be due to D3R and D2R agonism. It should be taken into consideration that change in correct response latency can also result from change in motivation and locomotor ability. To assess 5C-CPT performance various parameters should be considered concurrently (Van Enkhuizen et al., 2014). For example, if a change in correct latency without a concurrent change in correct magazine or responsivity index (motivation) is reported, it is highly likely that the change in correct latency is affected by decision making rather than non-cognitive disruptions such as motivation or motor disorders (Young et al., 2011; Van Enkhuizen et al., 2014). In this study, PD128907 did not change Go trial reward collection latency or responsivity index in a baseline-dependent manner, suggesting that the change in correct response latency observed in this study is likely to reflect a change in decision making instead of change in motivation. This finding suggests that D3 receptors makes an essential contribution to the regulation of decision-making (obsessions) and perseveration (compulsions) as measured by 5C-CPT. Rats with long latency to correct response at baseline (deficit in decision making) were more sensitive to the deleterious effect of PD 128907 on correct response latency. This finding is consistent with previous studies which report that D2/D3 receptor agonist quinpirole disrupts risk-based decision making of male rats as measured by probabilistic discounting task (St Onge et al., 2011). This study suggests that activation of D3 receptor by systemic administration of PD 128907 exert differential effect on decision making and perseveration as measured by 5C-CPT. Some rats displayed a massive increase in compulsivity or correct response latency while others were unaffected or even improved following administration of PD 128907. This study suggests that activation of D3 receptors potentiates compulsive behaviour in female rats that have a high perseveration score and a long latency to correct response at baseline (vulnerable population) compared to other rats. Thus, pharmacological manipulations of dopaminergic system specifically through activation of D3 receptors shows impairment in decision-making and compulsivity in rats characterized by deficit in decision making and high compulsivity.

Serotonergic and dopaminergic pathway has been linked with obsessive-compulsive disorder (OCD) (Stein, 2002). Quinpirole is a selective D2/D3 receptor agonist which is used to induce compulsive checking (repetitive behaviour) in male Lister hooded rats that matches the main symptoms of OCD in patients (Szechtman and Eliam 1998; Eagle et al., 2014). Animal models are essential for enhancing our understanding of the neurochemical basis of OCD and for preclinical testing of new therapeutic agents (Eagle et al., 2014). Many patients with OCD do not respond to current medications (Miodownik et al., 2015). The neural mechanism underlying OCD remains largely unknown. Obsessions which have been linked with cognitive dysfunction are difficult to demonstrate in animals. Repetitive rituals in OCD patients results from pathological doubt which has been linked to dysfunction in decision making process. Quinpirole induced compulsive checking (repetitive behaviour) in rats matches only part of compulsive behaviour in OCD patients (Szechtman et al., 1998; Eliam & Szechtman, 2005). The predictive validity of this model is not clear (Stuchlik et al., 2016). A better animal model is required to understand neural circuit implicated in OCD and to develop more effective treatments. 5C-CPT can be used to measure compulsivity (repetitive behaviour) and decision making at the same time (Tomlinson et al., 2014; 2015). PD 128907 induced perseverative responses resembles quinpirole induced compulsive checking like behaviour in an operant observing response task (Eagle et al., 2014). The present study suggests that PD 128907 effects on compulsive like behaviour and decision making in poor performing female lister hooded rats (rats with long correct response latency and high perseveration at baseline) in the 5C-CPT task matches two key features of OCD compulsions in humans (perseveration and indecision), suggesting that PD 128907 can be used to pharmacologically induce OCD in female rats. PD 128907 induced behavioural alterations in female lister hooded rats can serve as a better animal model of OCD. One way to determine predictive validity of PD 128907 induced compulsive like behaviour and indecision would be to test effects of existing OCD drugs such as clomipramine (tricyclic antidepressant) or Sulpiride which is a D2/D3 receptor antagonist in reversing these deficits (Eliam & Szechtman, 2005; Stein, 2002; Eagle et al., 2014). 5C-CPT is an effective tool for revealing detrimental effect of PD 128907 on compulsive behaviour and decision making. In addition, 5C-CPT can be used to measure compulsive behaviour in human and rat which may help in understanding the complex etiology of OCD (Young et al., 2013; Tomlinson et al., 2014). More work will be required to establish the predictive validity of this model.

In this study, group performance revealed no significant change in compulsive behaviour following administration of PD128907. Correlational analysis helped to reveal a relationship that would have otherwise gone unnoticed. To our knowledge, this is the first study to take advantage of the variation in individual baseline perseverative responses or correct response latency to demonstrate the presence of a rate-dependent relationship between baseline perseverative responses or correct response latency and the PD 128907-induced change. This finding is consistent with previous studies which report that the effect of pharmacological manipulation of dopaminergic system on inhibitory response control is dependent on heterogeneity of the performance prior to manipulation (Abdulkader et al., 2023a; 2023b). For example, selective inhibition of the dopamine transporter by administration of GBR12909 showed a tendency toward improving compulsive behaviour in a baseline dependent manner (Abdulkader et al., 2023a). Correlational analysis will provide the basis for understanding the neurobiology of disorders related to poor decision making and compulsivity such as OCD, gambling and addiction. The findings in this study, taken together with previous studies which report anti-gambling properties of D3 preferring antagonists, suggest that novel analogues of D3 receptor antagonists may have clinical utility for the treatment of compulsive behaviour related disorders such as OCD, compulsive drug seeking behaviour, substance abuse in subject with existing OCD, compulsive gambling and schizophrenia.

Behavioural results are clearly affected by strain, age and sex (Saré et al., 2021) It should be taken into consideration that Lister hooded rats are hyperactive, more impulsive and less attentive than Wistar rats (Jogamoto et al., 2020). This strain has higher vulnerability to develop compulsive like behaviour. Because of it is characteristics Lister hooded rats are commonly used in preclinical investigations of potential therapeutic agents for ADHD (Barnes et al., 2011; Tomlinson et al., 2014; Hayward et al., 2016). OCD and ADHD often coexist (Abramovitch et al., 2015). Male rats have been used in most animal models of OCD (Eliam & Szechtman, 2005; Stein, 2002; Eagle et al., 2014). However, OCD is more prevalent among females in adolescence and adulthood compared to males (Mathes et al., 2019). Gender differences in quinpirole sensitisation rat model for OCD have not been examined. Our results suggest the possibility of gender and strain dependent effect. Deficit in cognitive domains (attention, decision making and cognitive flexibility) may underlie the symptoms of OCD (Benzina et al., 2016). PD128907 has been reported to impair cognitive function in wild type mice (Watson et al., 2012). In the present study, Oldham’s method revealed a latent relationship between baseline accuracy and magnitude of change following administration of PD 128907 at the dose that affected perseverative correct responses and correct response latency, suggesting contribution of D3 receptors to attentional performance, compulsive behaviour and decision making in female rats. Deficit in attentional accuracy and decision making may have contributed to the compulsive behaviour in poor performing rats. This finding is consistent with previous studies which report that increases in perseveration and decreases in attentional accuracy in the 5-CSRTT sometimes co-occur (Grottick and Higgins, 2000; Greco et al., 2005; Le Pen et al., 2003). In the present study, a greater decrease in accuracy was observed in rats with low baseline accuracy (vulnerable population). This study is consistent with previous studies which report a significant decline in accuracy and increase in omission following systemic administration of PD 128907 (0.05 mg/kg) in mice version of 5-CSRTT (Zhu et al., 2017). The prefrontal cortex has a low D3 receptor density (Larson & Ariano, 1995). It has been reported that D3 receptors in the substantia nigra (SN) and ventral tegmental area (VTA) function as autoreceptors. Almost all dopaminergic neurons in SN and VTA express D3 receptors (Nakajima et al., 2013; Diaz et al., 2000). The impairment in selective attention observed in rats with low baseline accuracy could be due to the inhibitory effects of D3 receptor activation on mesocortical dopaminergic pathway which projects from VTA (ventral tegmental area) to the cortex (Mueller et al., 2017). Previous studies suggest that pro-cognitive and pro-social effects of D3 receptor antagonism could be due to indirect activation of cortical D1 receptor (Kiss et al., 2021). In addition, previous studies suggest that inhibition of D3 receptors upon introduction in frontal cortex, but not in nucleus accumbens, improved social recognition in rats (Loiseau et al., 2009). The improvement in social recognition was similar to effects of D1 selective agonist. On the other hand, D2 preferring antagonist had no effect (Loiseau et al., 2009). Furthermore, in the social recognition paradigm, the enhancement in performance following administration of D3 preferring antagonist (S33138) was accompanied by increase in fronto-cortical acetylcholine overflow (Millan et al., 2007), supporting the role of D3 receptors in modulating cholinergic neurotransmission (Watson et al., 2012). However, the mechanism by which D3 receptor inhibition results in increased ACh release is unclear. It may be argued that the impairment in accuracy observed after the lowest dose of PD 128907 (D3 receptor selective) was due to reduced cholinergic neurotransmission. This is the first study to report the presence of a latent rate-dependent effect following administration of a D3 receptor agonist which could help to translate preclinical findings into clinical studies (real life benefits). D3 preferring antagonists could be used as add-on therapy for treatment of OCD in patients.

In conclusion, these findings suggest that the concept of rate-dependency applies to effects of PD 128907 on compulsive behaviour, decision making and selective attention as measured by 5C-CPT. As expected, PD 128907 impaired compulsive behaviour and decision making in poor performing rats (vulnerable population). A correlative analysis (e.g., Oldham’s method) may be particularly effective in revealing drug effects on behaviour. PD 128907 induced behavioural alterations in female lister hooded rats can be used as an animal model of OCD which may facilitate the discovery of new therapeutic agents for OCD. Future work can expand the current study, for example, 5C-CPT has been reverse translated to human, enabling compulsive behaviour to be studied in both OCD patients and rodents.

## Declaration of conflicting interests

The author(s) declared no potential conflicts of interest with respect to the research, authorship, and/or publication of this article.

## Funding

The author(s) received no financial support for the research, authorship, and/or publication of this article.

## Notes

### Competing Interest Statement

The authors have declared no competing interest.

